# Only the fastest corticospinal fibers contribute to beta corticomuscular coherence

**DOI:** 10.1101/2020.11.18.387282

**Authors:** J. Ibáñez, A. Del Vecchio, J. C. Rothwell, S. N. Baker, D. Farina

## Abstract

A common way to study human corticospinal transmission is with transcranial magnetic stimulation. However, this is biased to activity in the fastest conducting axons. It is unclear whether conclusions obtained in this context are representative of volitional activity in mild-to-moderate contractions. A possible alternative to overcome this limitation is to study the corticospinal transmission of endogenously generated brain activity. Here we study the transmission speeds of cortical beta rhythms travelling to the muscles during steady contractions. To do this, we introduce new methods to improve delay estimates in the corticomuscular transmission of beta rhythms, and which we validate both in simulations and experimentally. Applying these approaches to experimental data from humans, we show that corticomuscular beta transmission delays are only 1-2ms longer than expected from the fastest corticospinal pathway. Simulations using realistic distributions of the conduction velocities for descending axons projecting to lower motoneurons suggest two scenarios that can explain these results: either a very small fraction of only the fastest corticospinal axons selectively transmit beta activity, or else the entire pool does. The implications that these two scenarios have for our understanding of corticomuscular interactions are discussed in the final part of this manuscript.

**SIGNIFICANCE:** We present and validate an improved methodology to measure the delay in the transmission of cortical beta activity to tonically active muscles. The estimated corticomuscular beta transmission delays which this yields are remarkably similar to those expected from transmission in the fastest corticospinal axons. A simulation of beta transmission along a pool of corticospinal axons using a realistic distribution of fiber diameters suggests two possible mechanisms by which fast corticomuscular transmission is achieved: either a very small fraction of descending axons transmits beta activity to the muscles or, alternatively, the entire population does and natural cancellation of slow channels occurs due to the distribution of axon diameters in the corticospinal tract.

## Introduction

The corticospinal tract plays a fundamental role in dexterous movements in primates (Lemon, 2008). To date, studies of corticospinal function have relied on microelectrode recording or stimulation in animals, and non-invasive stimulation in humans (Rothwell, 2007, 2012; Lemon, 2008). Both of these approaches are biased towards the largest cells with the fastest conducting axons; by contrast little is known of the contribution of slower fibers, even though these make up the overwhelming majority of the corticospinal tract (Kraskov et al., 2019). One way to overcome this limitation might be to study the transmission of endogenous motor commands from cortex to muscle, since the different corticospinal axon calibers will impose their own characteristic conduction times on the signal propagation.

A particularly advantageous state to examine slow and fast corticospinal transmission to motoneurons might be during voluntary tonic contractions, when motor cortical circuits generate oscillatory activity in the beta band (15-30 Hz) (Baker et al., 1997). These oscillations are known to be carried by fast corticospinal cells (Baker et al., 2003), and to synchronize motoneurons (Williams and Baker, 2009; Negro and Farina, 2011) leading to significant cortico-muscular coherence (Conway et al., 1995; Baker et al., 1997; Salenius et al., 1997). Reliable characterization of beta time delays between cortex and muscles could quantify how oscillations are carried by the different conduction velocity divisions of the corticospinal tract.

Many previous studies have studied the delay between coherent cortical and muscle beta activity (Brown et al., 1998; Halliday et al., 1998; Gross et al., 2000; Mima et al., 2000; Riddle and Baker, 2005a; Govindan et al., 2006). Determining oscillatory delays is relatively straightforward if there is transmission in only one direction (Halliday et al., 1995). However, the motor cortex not only sends oscillations down to the spinal cord (Baker et al., 2003), but also receives ascending oscillatory input over afferent pathways (Baker et al., 2006; Baker, 2007). Such bidirectional coupling can generate erroneous estimates using simple correlational measures like coherence (Cassidy and Brown, 2003; Witham and Baker, 2012; Campfens et al., 2013). In modelled data, delay estimates closer to the known ‘ground truth’ are generated by approaches such as Granger Causality (directed coherence) (Witham et al., 2010). When applied to physiological recordings, directed coherence measures longer delays than those expected from the fastest corticospinal axons, suggesting that both fast and slow axons conduct the beta oscillations (Witham et al., 2010, 2011). However, there is considerable inter-subject variability – a factor of approximately twofold between the slowest and fastest delay estimates (Witham et al., 2011). The extent to which slow fibers participate in signal transmission is a fundamental property of the motor system; it seems implausible that it should exhibit such profound biological variation, which might instead be generated artifactually by methodological issues.

Here we study the contribution of fast and slow corticospinal fibers to corticomuscular beta transmission. To do this, we first propose and validate two critical improvements to measuring corticomuscular delays. Firstly, we previously showed that directed coherence extracts accurate delay estimates using simulated data with a broad range of spectral frequencies. However, corticomuscular beta coherence only involves transmission of signals within a relatively narrow band. Surprisingly, here we find that this can impact delay measures, and that an alternative approach in the time domain (cumulant density, (Halliday et al., 1995)) is more reliable than directed coherence. Secondly, past work used the interference electromyogram (EMG) to represent peripheral activity; this will incorporate a dependence on the shape of motor unit action potential waveforms (Williams and Baker, 2009), which is a source of spurious variability. Here we instead used the spiking activity of pools of motor units blindly extracted from surface recordings (Farina and Holobar, 2016). The resulting improved corticomuscular transmission delay estimates are compared with conduction times for fast corticospinal transmission determined by noninvasive brain stimulation. Finally, we use a model to understand the implications of our results for passage of motor commands over both fast and slow corticospinal axons.

## Methods

This work presents data from three experiments. Experiment 1 involves simulations of bidirectional beta transmission between two sources (Fig. 1A). The aim of this experiment was to test the reliability of the directed coherence and the cumulant density function to estimate temporal delays between coherent band-limited signals.

**Figure 1.**
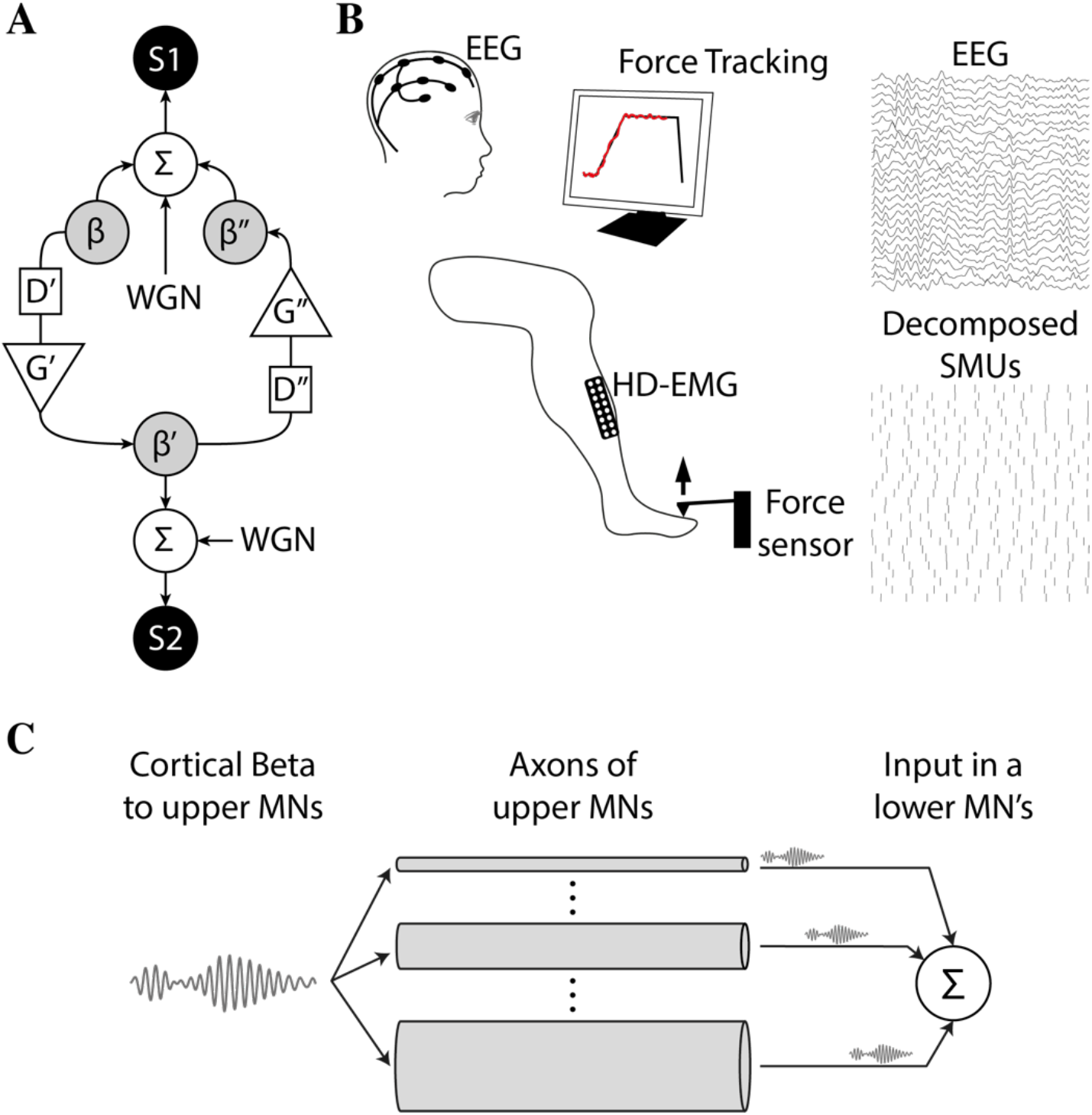
The study presented here is based on data from three experiments. (A) Schematic of the simulation used in Experiment 1 to test the reliability of the cumulant density and directed coherence methods to estimate interaction delays in a bidirectional system in which only a limited band of frequencies is transmitted between sources S1 (simulating a cortical source) and S2 (simulating a muscle source). WGN stands for white gaussian noise; ∑ indicates the summation of input signals; D’/D’’ and G’/G’’ represent the delays and gains in the S1→S2 and S2→S1 directions. Delays were generated with band-pass filters that restricted the transmitted signal (ß/ß’/ß’’) to a specific frequency band centered around 21 Hz. (B) Schematic representation of the set-up in the main recording in Experiment 2. Subjects sat with their right foot placed against a strain gauge measuring forces produced by dorsiflexing the ankle. During the recording blocks, subjects were asked to track a ramp-and-hold force trace on a screen by producing torques with their ankle (hold part at 10 % of the maximum voluntary contraction level). High-density EMG (HD-EMG) and EEG were recorded and offline processed to study the characteristics of the interactions between scalp signals and the composite spike train resulting from combining the information of all the identified SMUs from the TA. M1→muscle beta transmission delay estimates were the main outcome measure. (C) Simulation run in Experiment 3 to analyze the conditions that can explain the observed beta transmission delays in Experiment 2, in the light of what is known about the distribution of axon diameters in upper motoneurons projecting to the motoneuron pool.

In Experiment 2, electroencephalography (EEG) and high-density electromyography (HD-EMG) of the tibialis anterior muscle (TA) were acquired concurrently in human subjects during sustained ankle dorsiflexions (Fig. 1B). Additionally, in a subset of subjects, motor-evoked potentials (MEPs) induced by transcranial magnetic stimulation (TMS) were measured to estimate the minimum corticomuscular transmission delays (Day et al., 1989). Recordings in Experiment 2 are used for three purposes. First, they are used to confirm the results obtained by simulation in Experiment 1 regarding the comparison between beta transmission delay estimates using the directed coherence and the cumulant density function. Second, recordings are used to compare corticomuscular beta transmission delays obtained using single motor unit (SMU) activity and the interference EMG obtained using standard surface montages. This allows us to determine the influence of the shapes of motor unit action potentials (MUAPs) on beta transmission delay estimates. Finally, data from Experiment 2 are also used to test the relation between beta transmission delays and subjects’ heights, and to compare transmission delays with MEP latencies to infer the conduction velocity of the descending pathways transmitting beta activity to the muscles.

Finally, in Experiment 3, we simulate the propagation of beta activity along the axons of upper motoneurons with realistic distributions of axon diameters (and conduction velocities), and the subsequent summation of the delayed signals in the soma of lower motoneurons of a muscle (Fig. 1C). We use the results from this analysis to infer, in the light of the results from Experiment 2, how beta may be transmitted to the muscles by populations of upper motoneurons.

### Experiment 1: Simulation of a bidirectional system transmitting band-limited signals

We simulated a bidirectional system with two sources (S1 and S2) generated by combining independent white Gaussian noise with a shared beta signal (ß) constrained in a specific frequency band around 21 Hz (Fig. 1A). Simulations were meant to resemble bidirectional corticomuscular interactions that are typically constrained within the beta band (Baker 2007), and used a sampling rate of 256 Hz and a duration of 120 s. To simulate cortical beta projections to muscles, ß was transmitted from S1 to S2 (ß’) with a delay (D’) of 19.5 ms and a gain (G’) of 0.2. Then, to simulate afferent projections to the brain, ß’ was transmitted from S1 to S2 (ß’’) with a delay (D’’) of 31.3 ms and a gain (G’’) of 0.2. The delays here were defined to make the simulated system comparable to previous similar works (Witham et al., 2010). Gains were empirically chosen to obtain coherence results similar to those obtained with experimental data from humans (e.g., results obtained in Experiment 2 here). To understand the influence of the bandwidth of the beta signals, we ran different simulations with ß signals being band-pass filtered within frequency bandwidths between 4 Hz and 40 Hz, in incremental steps of 2 Hz, and centered around 20-22 Hz (Kilner et al., 1999). Signal-to-noise ratios (SNR) of the beta contributions to S1 and S2 between 0 dB and 12 dB (in steps of 1 dB) were tested for each bandwidth considered. All the parameters in the model were determined empirically to make the results obtained from simulations match the average results with real human data (*i.e.*, results obtained in Experiment 2).

### Experiment 2: Recordings of EEG and HD-EMG during tonic contractions of the TA

Nineteen healthy subjects (2 females; age range 23 - 35 years) were recruited. The average height was 178 ± 6 cm (range 165-187 cm). All subjects gave their informed consent before participating in the study. The study was approved by the University College London Ethics Committee (Ethics Application 10037/001) and conducted in accordance with the Declaration of Helsinki.

#### Recordings

At the beginning of each session, subjects were asked to sit on a straight chair with their knees flexed at a 90° angle and their right foot placed beneath a lever measuring ankle dorsiflexion forces during isometric contractions (Fig. 1B). The ankle forces together with high-density EMG (HD-EMG) recordings of the TA were acquired by a multichannel EMG amplifier (Quattrocento, OT Bioelettronica, Torino, Italy). HD-EMG grids (5 columns and 13 rows; gold coated; 1-mm diameter; 8-mm interelectrode distance) recorded the myoelectrical activity of the TA. Before placing the grids, the skin was shaved and cleansed with a 70 % alcohol solution. HD-EMG signals were acquired in monopolar mode.

EEG was recorded with an ActiChamp amplifier (polarity defined as negative upwards; BrainProducts, Munich, Germany) from 63 Ag/AgCl scalp electrodes using a 10/10 standard layout. Recordings were referenced to POz and the ground electrode was placed on AFz. Electrode impedances were maintained below 10 kOhms. A sampling frequency of 1000 Hz was used to acquire the data.

A common digital signal was used to synchronize HD-EMG and EEG data offline.

#### Experimental paradigm

After a standardized warm up, subjects performed three maximal voluntary isometric ankle-dorsiflexion contractions (MVC) that were separated by at least 30 s. During the MVCs, subjects were instructed to contract as much as possible for at least 3 s. The peak force value across the MVCs corresponded to the final MVC value and was digitally recorded. The MVC across subjects was 180 ± 82 N. For the main recordings, subjects were asked to follow ramp-and-hold force trajectories displayed on a monitor. The trajectories consisted of 2-s ramp contractions from 0 % to 10 % MVC followed by a 60-s holding phase at 10 % MVC. During the holding phase, subjects were instructed to keep an isometric force of 10 % MVC. The ramp-and-hold contractions were repeated twice by each subject, with a 2 min rest in between.

#### Recording of MEP latencies

Five subjects recruited for Experiment 2 (average height 179 ± 4 cm) took part in a second experiment performed on a different day, which measured the latencies of MEPs in the TA following TMS over M1. For this experiment, bipolar EMG from the TA was recorded with two surface Ag-AgCl electrodes 2 cm apart (WhiteSensor 40713, Ambu). The ground electrode was placed on the right ankle. EMG signals were amplified, band-pass filtered between (20-2000 kHz, Digitimer D360 amplifier, Digitimer Ltd, United Kingdom) and acquired at 5 kHz sampling rate with a data acquisition board (CED Power1401, Cambridge Electronic Design Ltd, UK) connected to a PC and controlled with Signal V6 software (also CED). A standard TMS device connected to a 110 mm double-cone coil (Magstim 200^2^, The Magstim Company, UK) was used to stimulate the TA representation of the left M1. The coil was held tangentially on the scalp at an angle of 0° to the mid-sagittal plane to induce a posterior–anterior current across the central sulcus. The motor hotspot was determined by searching for the position where consistent MEPs could be seen in the TA at rest. Active motor threshold (AMT) was defined as the lowest intensity to evoke a MEP of at least 0.2 mV peak-to-peak amplitude in 5 of 10 consecutive trials while subjects gently contracted their muscles (around 0.15 mV peak-to-peak background activation). Thereafter, 15 TMS pulses with an intensity of 1.4x AMT were delivered while subjects maintained contraction. This gave clear MEPs in the TA. In offline analysis, MEP onset latency was determined via visual inspection on a trial by trial basis from the raw EMG traces and the average latency was determined (Rocchi et al., 2018; Ibáñez et al., 2020).

### Experiment 3: Simulation of corticospinal beta transmission to lower motor neurons through multiple axons with different conduction velocities

We simulated the transmission of beta activity along the descending axons of upper motoneurons directly innervating cells in the motoneuron pool of a muscle. Specifically, we simulated how cortical beta inputs would be summated at the soma of lower motoneurons when considering a distribution of transmission delays across the descending axons of the upper neurons based on previously published results (Firmin et al., 2014). Therefore, we used the list of observed axon diameters from Firmin’s paper (monkey CS28; raw data kindly provided by R.N. Lemon) and converted them into axon transmission delays using a Hursh factor of 6. In our tests here, we assumed axon lengths from 400 mm to 1000 mm. Previous reports give estimated central motor conduction time for the TA as ~14.5 ms (Jaiser et al., 2015). Allowing 3-4 ms for transsynaptic depolarizations at the cortical and spinal levels leaves ~11 ms for axonal conduction. Considering that the fastest corticospinal conduction velocity is 76.2 m/s (Firmin et al., 2014) gives an axon length of 840 mm. Our simulated length range therefore includes the best estimate of the actual length relevant for the TA muscle. A lognormal function was fitted to the transmission delays calculated from Firmin et al’s data; a Kolmogorov-Smirnov test failed to reject the null hypothesis that the experimental data comes from a lognormal distribution (P = 0.056). Our model considered 10^6^ descending axons. The beta input to the descending axons was modelled as 20 s of white Gaussian noise band-pass filtered in the beta band (Butterworth, 3rd order, band 20-30 Hz). Delays according to the distribution of descending axons were applied to the simulated beta input in the frequency domain (McGill and Dorfman, 1984). Finally, all the delayed versions of the simulated beta input were summed and the group delay between the resulting signal and the input was estimated from the phase-frequency regression in the coherence function (obtained using the periodogram method with 1-s segments, and discarding the first 1-s segment containing a transitory period). Note that this way of estimating delays is the most suitable one for unidirectional transmission of sinusoidal signals (Rosenberg et al., 1989). Simulations were performed using a sampling rate of 256 Hz.

The above simulation was run first by simulating the transmission of the input beta signal through all the axons (100% of the axons) generated by the fitted lognormal distribution. Then, the same simulation was repeated after applying a set of thresholds to the axon diameters that could transmit beta rhythms. Each threshold was used as a lower bound applied to axon diameters, so that only a certain percentage of the fastest axons (with diameters above a given threshold) contributed to beta transmission. The following percentages were tested: 0.1%, 0.5%, 1%, 5%, 10%, 25%, 50%, and 50%.

Finally, in addition to group delays, normalized amplitudes of the resulting signals after combining all the delayed versions of the simulated beta input were estimated to measure the strength of the transmitted information. This was done by obtaining the root-mean-square (rms) amplitude of the signals resulting from summing the contributions of all the axons transmitting the beta input, and dividing the resulting value by the total number of axons considered (10^6^) and by the rms amplitude of the simulated input (as with the delay estimates, the first 1-s segment was ignored here). These amplitudes thus depended on the amount of cancellation among the channels transmitting beta activity and on the number of axons included by each diameter threshold.

### Data analysis

#### HD-EMG decomposition into motor unit spike trains

The main results in Experiment 2 were obtained using spiking activity of SMUs instead of the interference EMG. In offline analysis, the HD-EMG signals were first band-pass filtered (20-500 Hz band, 2nd order zero-lag Butterworth filter) and subsequently decomposed into SMU spike trains by blind source separation techniques described in previous works (Holobar et al., 2014; Negro et al., 2016). The accuracy of the decomposition algorithm used has been previously demonstrated both by simulation and experiment over the full recruitment range of the TA (Farina and Holobar, 2016; Del Vecchio and Farina, 2019). SMUs recruited more than 10 s after subjects reached the 10 % MVC level in each recorded block and SMUs with a standard deviation in instantaneous firing rate higher than 20 Hz were discarded. The remaining SMUs were used to construct a composite spike train to be used in subsequent connectivity analyses. This was done by summing the binary activity of all the trains of pulses from the valid SMUs. The spike time of each SMU corresponded to the onset of its MUAP. To estimate this onset, double differential signals were obtained from the monopolar HD-EMG recordings along the columns of the electrodes grid (*i.e.*, approximately along the longitudinal direction of muscle fibers). Then MUAPs were obtained via spike-triggered averaging (Farina and Holobar, 2016) and the channel in the recording grid showing the largest ratio between the peak-to-peak amplitudes of the MUAP and preceding noise (in the interval [−40, −20] ms before the peak time of the MUAP) was selected to estimate the MUAP onset time. Using this channel, the time at which the average MUAP first crossed a threshold set at five standard deviations above the mean basal activity was defined as the MUAP onset time. Onset estimates were checked visually, and any errors from this automated approach corrected manually.

#### Derivation of monopolar and bipolar EMG from HD-EMG recordings

To study how MUAPs affect corticomuscular delay estimates, we compared EEG→SMU and EEG→EMG beta transmission delays estimates. Bipolar and single channel monopolar recordings were derived from the originally recorded monopolar HD-EMG signals in order to simulate the recordings that would have been obtained using typical EMG montages. To derive the bipolar EMG, two sets of five neighboring electrodes in the electrodes grid (with the central electrode of the grid corresponding to the belly of the muscle) were averaged, the difference of these averages was then computed to derive EMG signals recorded by large (standard) electrodes having an interelectrode distance of 1.6 cm (Del Vecchio et al., 2017). Single channel monopolar EMG was obtained by averaging the recordings of the five central positions in the recording grid. Both bipolar and monopolar EMG derivations were then analyzed both with and without rectification (Negro et al., 2015).

#### EMG synthesis using motor unit activity

To study if differences between EEG→SMU and EEG→EMG beta transmission delay estimates were only due to the filtering effect of MUAPs, we synthetized EMG signals from the decomposed SMU activity and compared the estimates of beta delays with the ones obtained using the real EMG (Del Vecchio and Farina, 2019). For this purpose, synthesized signals were obtained from the identified SMU activity by convolving the spiking activity with the estimated shapes of MUAPs (i.e., using the MUAPs shapes as filters). These shapes were obtained using spike-triggered averaging (Farina and Holobar, 2016). The used window to extract peri-spike segments had the following expression:

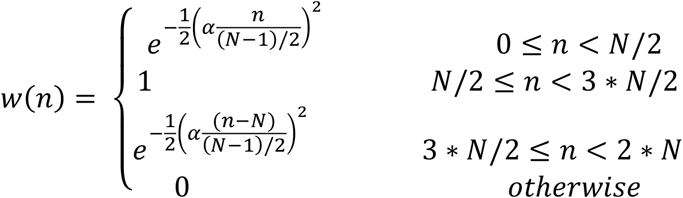

where N is the length of the central flat part of the window (set to 50 ms), n is the sample index and α is a constant set to 2.5. This window tapered the extracted MUAP shapes at the onset and offset so that the estimated MUAP shapes used as filters to synthetize EMG activity did not present any sharp transitions. Individual MUAP shapes were extracted by multiplying *w(n)* with EMG segments around the estimated spike times. The MUAP shape of for each SMU was then obtained by averaging all the individual MUAPs of the corresponding unit.

#### EEG pre-processing

EEGLAB toolbox for Matlab was used for the EEG pre-processing (Delorme and Makeig, 2004). First, EEG signals were bandpass filtered between 0.5-45 Hz (Butterworth IIR filter, 3rd order). An independent component analysis algorithm (BINICA) was used to visually identify and remove artefacts due to electrode noise, muscle activity, eyeblinks and eye movements. Laplacian derivations (subtracting the average activity of the closest equidistant channels) were computed for electrodes FCz, C1, Cz, C2 and CPz. The resulting signals from these derivations were used for the connectivity analysis described below. Specifically, the channel with which corticomuscular coherence amplitude was highest in the beta band in each case was selected for further analysis (Ibáñez et al., 2014). The channel selected was Cz in most cases (16/ 19 subjects in the analysis using SMUs) (Petersen et al., 2012).

#### Connectivity analysis

A key part of our study is the analysis of the connectivity between continuous (EEG) and spiking (SMUs) signals. The main results presented here were obtained using the previously validated Neurospec 2.11 toolbox for Matlab (Mathwoks Inc., USA) aimed to analyze nonparametric directional connectivity between time series and point process data (www.neurospec.org) (Halliday, 2015). In brief, this framework uses a combined time and frequency domain approach to decompose the standard coherence function by direction. Here, this directional coherence estimation method is used to extract, from the conventional EEG-SMU (or EEG-EMG) coherence function, the part resulting from cortical projections to muscles (EEG→SMU or EEG→EMG coherence). For the identification of significant coherences, we use Neurospec’s estimates for the upper 95% confidence limit (Rosenberg et al., 1989), defined as 1 − 0.05^1/(*L*−1)^, where L is the number of segments from which the Fourier transform is computed. In all cases, coherence analysis with Neurospec was done using segments of 1 s and multitapers (three tapers) for spectral estimation.

The Neurospec 2.11 framework also allows the characterization of corticomuscular interactions in the time domain by estimating the cumulant density function (cross-covariance) together with upper and lower 95 % confidence limits to determine significant peaks based on the assumption of uncorrelated processes. These confidence limits are set as 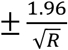, where R is the number of data points considered. The cumulant density function and the confidence limits are used here to estimate the lag with which the possible correlation existing between EEG and muscle activity is strongest and to assess its significance. To focus on the delays of corticomuscular transmissions descending from the brain (EEG→SMU/EMG transmission delays), the time point with the largest peak found at positive lags was determined. Peak time was used because EEG recordings were performed with an inverted polarity (negative upward), implying that positive deflections relate to periods during which dendritic trees of pyramidal cells in the motor cortex receive predominating excitatory input (Baker et al., 2003; Farmer et al., 2004; Zrenner et al., 2017).

Directed coherence (spectral Granger causality) was also used to replicate results obtained in previous studies of transmission delays in corticomuscular beta interactions. Specifically, we used the same implementation of directed coherence as in Witham et al. (2010, 2011). In brief, the studied signals were down-sampled to 512 Hz and autoregressive models of order 256 were fitted to blocks of 10 s, followed by averaging of model coefficients (Witham et al., 2010). Geweke’s normalization of directed coherence was used (Geweke, 1982). The 95 % confidence intervals for significance assessment were obtained using Monte Carlo simulation as in previous studies (Witham et al., 2010, 2011). Beta transmission (group) delays were determined from the slopes of the linear functions fitted to phase values corresponding to frequencies at which directed coherence was significant in the range between 12 Hz and 30 Hz. Delay estimates were only considered reliable if four or more phase points could be used for linear fitting.

Finally, in the analysis of the simulated data in Experiment 1, apart from using the cumulant density function and the directed coherence, beta transmission delays were also estimated using the phase of the spectral coherence (obtained using the coherence estimate calculated using Neurospec211 toolbox as explained above). This provided results that could be compared with previous simulation studies (Witham et al., 2010).

#### Statistics

Statistical analyses of results in Experiment 2 were performed using SPSS (version 22.0; IBM, USA). All results are reported as the mean ± SD (unless specified otherwise) and considered significant if P < 0.05. Normality of data distribution was assessed using Lilliefors tests. Comparison of means and variances of EEG→SMUs delay estimates based on the cumulant density function and the directed coherence was done using a paired t-test and an F-test. Paired t-tests were run between delay estimates obtained with SMU activity and with each of the considered EMG recording configurations (monopolar or bipolar; rectified or unrectified). Pearson’s correlation (one-tailed) was used to study the possible relationship between corticomuscular beta transmission delay estimates and subjects’ heights. Finally, Pearson’s correlation (two-tailed) was used to measure the correlation between delay estimates obtained using real EMG and the EMG synthesized from the spiking trains of the decomposed SMUs as explained in previous sections. For this analysis, we rejected data points which were two standard deviations above or below the mean delays combining the results from real and synthesized data. This led to the removal of five points before testing the correlation, and it was done to disregard possible wrong delay estimates from the cumulant density function either using the real or synthetized EMG (due to spurious peaks in the cumulant density; see for example the extreme values with the monopolar unrectified configuration in the results section).

## Results

This section is divided into two sections. In the first, results from Experiments 1 and 2 are used to indicate several factors that need to be considered to estimate corticomuscular beta transmission delays reliably. The second section then provides reliable estimates of the beta transmission conduction velocities. This, combined with results from Experiment 3, allows us to infer which fibers of the corticospinal tract contribute to corticomuscular beta transmission.

### PART I - Reliably estimating beta corticomuscular transmission delays

#### PART I-A. The cumulant density provides more reliable corticomuscular beta transmission delay estimates than the directed coherence

In this section, we first use simulations of a bidirectional system transmitting signals with different bandwidths and signal to noise levels to explore the measurement of beta transmission delays by different methods. Then, we repeat this comparison with experimental data from healthy subjects.

Figure 2A shows the total coherence (black lines) and the directed coherence in the S1→S2 direction (grey shaded areas) for representative simulations with different bandwidth and SNR levels. Transmission delays were estimated using three methods. First, delays were obtained from a regression line fitted to the phase of the standard coherence function, which is a common approach in studies of corticomuscular coherence (Pogosyan et al., 2009; Petersen et al., 2012). Estimates in this case were in the range [− 0.1, +0.1] ms for all bandwidths and SNR levels tested, representing a consistent underestimate of the true value by about 20 ms (results not plotted in a figure). This is in line with previous work: the coherence spectrum does not estimate delay well in systems with bidirectional interactions (Witham et al., 2010).

**Figure 2.**
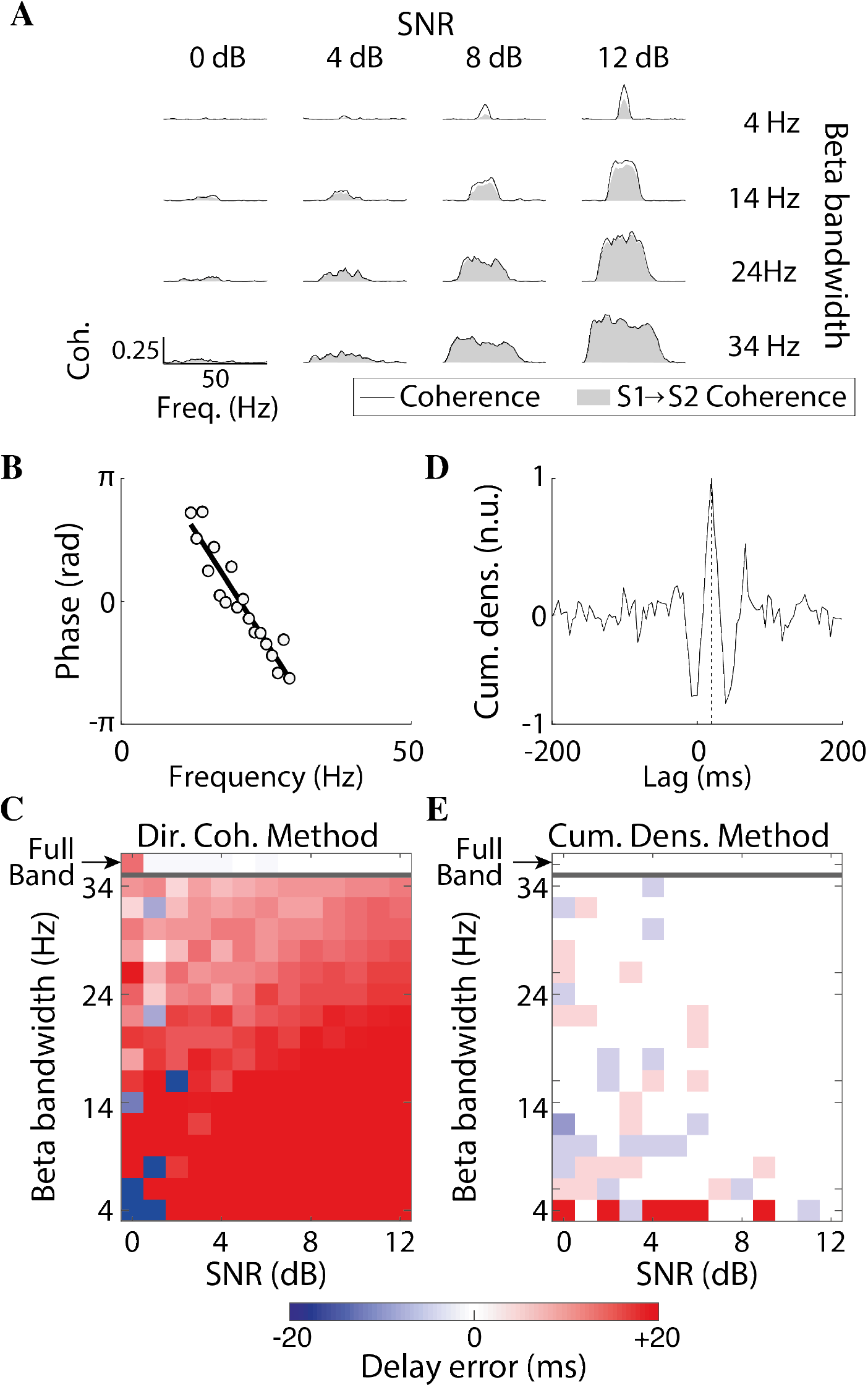
Beta transmission delay estimates using the directed coherence and the cumulant density function with data from Experiment 1. (A) Coherence (black lines) and S1→S2 coherence (grey areas) for different simulated conditions. From top to bottom rows, results for different beta bandwidths of the transmitted signals are plotted: 4 Hz, 14 Hz, 24 Hz, and 34 Hz. From left to right column, results for different SNRs of the transmitted signals: 0 dB, 4 dB, 8 dB and 12 dB. (B) Example of linear regression of the directed coherence phase–frequency relationship in the S1→S2 direction used to estimate the beta transmission (group) delay. (C) Errors in the transmission delay estimates in the S1→S2 direction based on the directed coherence. (D) Example of beta transmission delay estimation based on the cumulant density. The vertical dashed line indicates the transmission delay with which the maximum peak is found in the cumulant density. (E) Errors in the transmission delay estimates in the S1→S2 direction based on the cumulant density.

The second method tested was directed coherence (Witham et al., 2011; Witham and Baker, 2012). An example (for 18 Hz bandwidth, SNR 2 dB) of a linear fit to the phase-frequency spectrum for directed coherence is shown in Fig. 2B. Figure 2C shows the error in delay estimate (relative to the known ‘ground truth’ of the simulation), for varying levels of bandwidth and SNR. When the transmitted signal covered the full spectrum of frequencies (white noise), delay estimates based on the directed coherence were accurate for SNR> 0dB (white shading, top row Fig. 2C), as reported by previous work (Witham et al., 2010). However, when the transmitted signals had more limited bandwidth, estimates at high SNR were biased towards delays longer than the true values. Measurements for SNR = 0 dB were variable and unreliable.

Finally, the cumulant density was used to estimate delay in the time domain, by finding the latency of the positive peak (see example in Fig. 2D). These delays estimates were accurate for a wide range of parameters, and only showed appreciable errors for bandwidths of 4 Hz (Fig. 2E). This lack of reliability for narrow band signals is due to the large oscillations appearing in the cumulant density in these cases, which has been previously noted to introduce errors (Halliday, 2015).

Experimentally-recorded measurements from Experiment 2 were analyzed to obtain beta transmission delay estimates or comparison with the simulations in Experiment 1. To obtain reliable corticomuscular transmission delays, spiking activity of pools of SMUs was used. On average, 18 ± 8 SMUs were extracted per subject from HD-EMG (range 5:29 SMUs). The average discharge rate across all SMUs was 11.0 ± 1.6 pps.

Significant EEG-SMUs coherence in the beta band was observed in 18 out of the 19 subjects tested. The cortico-muscular coherence peak was 0.09 ± 0.08, compared to the limit used to define significant coherence peaks of 0.01 ± 0.00. Figure 3A shows the individual and average coherence estimates in the descending direction (EEG→SMUs). The average peak amplitude of the EEG→SMUs coherence was 0.07 ± 0.06, and it was significant in 16 cases. We did not find clear coherence peaks at other frequencies outside the beta band across subjects. In addition, coherence in the SMUs→ EEG direction was smaller and variable (on average it was 0.02 ± 0.02). These results indicate that, in the context of sustained mild contractions with the TA, beta is the only frequency band within which rhythmic cortical signals are consistently transmitted to the TA, and that the contribution of descending projections to the total coherence is stronger than the contribution of ascending projections.

**Figure 3.**
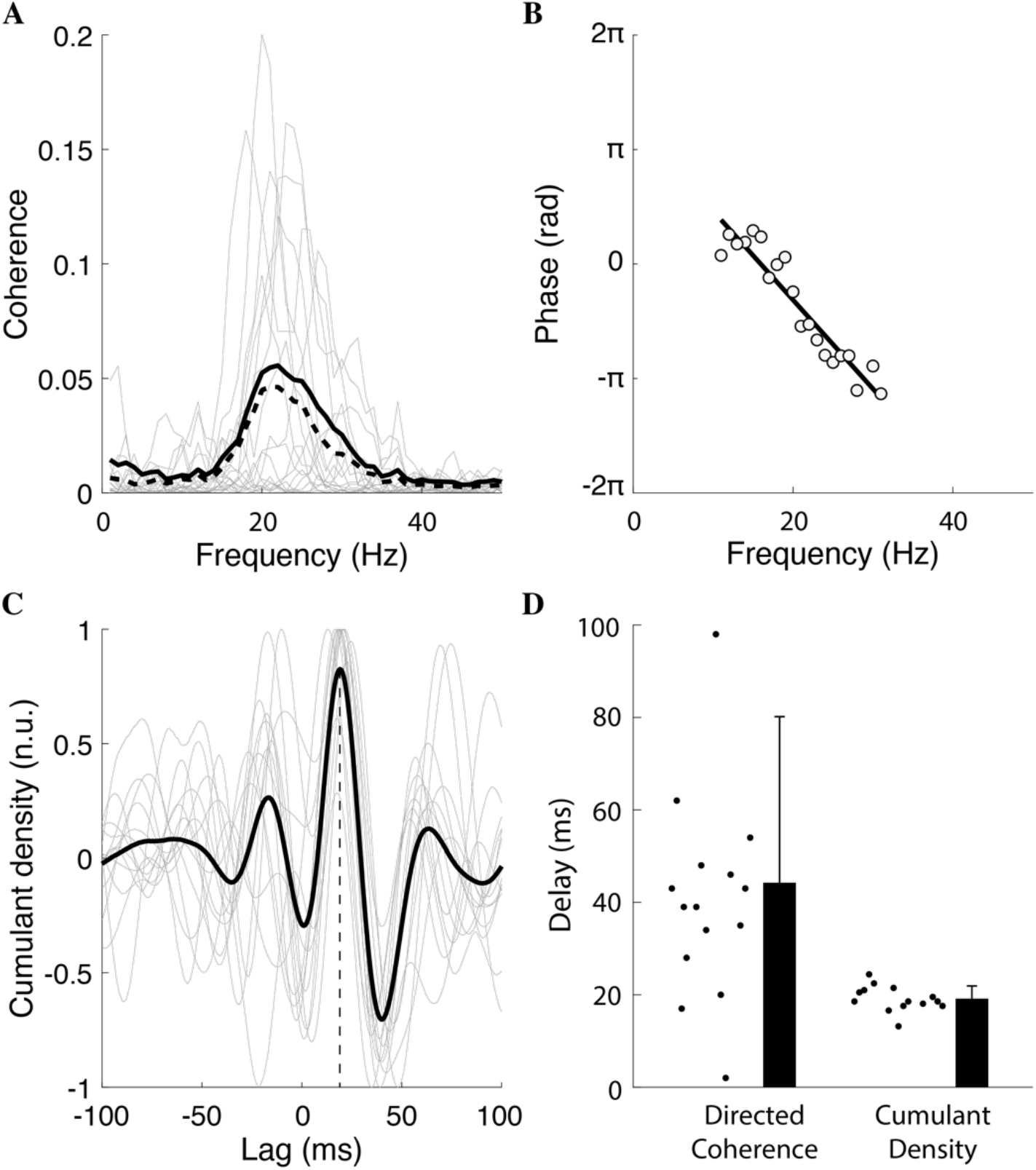
Beta transmission delay estimates using the directed coherence and the cumulant density function using data from Part 2. (A) Average coherence (solid black), EEG→SMU coherence (dashed black) and single-subject EEG→SMU coherence (grey). (B) Example of the phase spectrum from the directed coherence in the EEG→SMU direction to estimate transmission delays. Data from one subject (white circles) and linear fit to estimate the delay (black line). (C) Average (black) and single-subject (grey) cumulant densities. A vertical dashed line is used to indicate the lag at which the average peak in the cumulant density is found. (D) Beta EEG→SMU transmission delay estimates using the cumulant density function and the directed coherence. Grey circles represent individual results.

Figure 3B shows the linear fit to the EEG→SMUs directed coherence phase for a representative subject; the slope indicated a 38.8 ms delay for this example. The average cumulant density (Fig. 3C) showed a positive peak for most individual traces at a positive time lag. Significant cumulant density peaks at positive lags were seen in 14 subjects. Using the directed coherence, significant delay estimates could be obtained from 15 subjects. Average EEG→SMUs delays based on these two methods were 19.1 ± 2.7 ms and 44.2 ± 35.9 ms, respectively (Fig. 3D). These differences are in line with those from simulated data in Experiment 1. In the group of subjects from whom significant EEG→SMUs delay estimates could be obtained using both, the cumulant density and directed coherence, the mean and variance of delays were significantly larger with the directed coherence than with the cumulant densities (t (12) = −3.556, p = 0.004; F(1,12) = 12.645, p = 0.004).

#### PART I-B. Pooled SMU activity gives better delay estimates than EMG

Surface EMG signals are the result of convolving spike trains with the corresponding motor unit action potentials (MUAPs) at the skin surface (Farina and Merletti, 2001). To study the influence of the filtering effect of MUAPs on descending spike trains, EEG→SMUs beta transmission delay estimates were compared with those obtained using monopolar and bipolar EMG signals with and without rectification using the data from Experiment 2 (Fig. 4A). Overall, delay estimates based on SMUs (19.1 ± 2.7 ms; average of 14 subjects with significant delay estimates) were significantly lower (P < 0.001) than with any EMG-based configuration: 25.1 ± 4.6 ms using bipolar EMG and rectification (12 subjects with significant estimates); 36.1 ± 8.7 ms using bipolar EMG and no rectification (10 subjects with significant estimates); 24.6 ± 3.2 ms using monopolar EMG and rectification (14 subjects with significant delay estimates); 38.9 ± 18.7 ms using monopolar EMG and no rectification (17 subjects with significant estimates).

**Figure 4.**
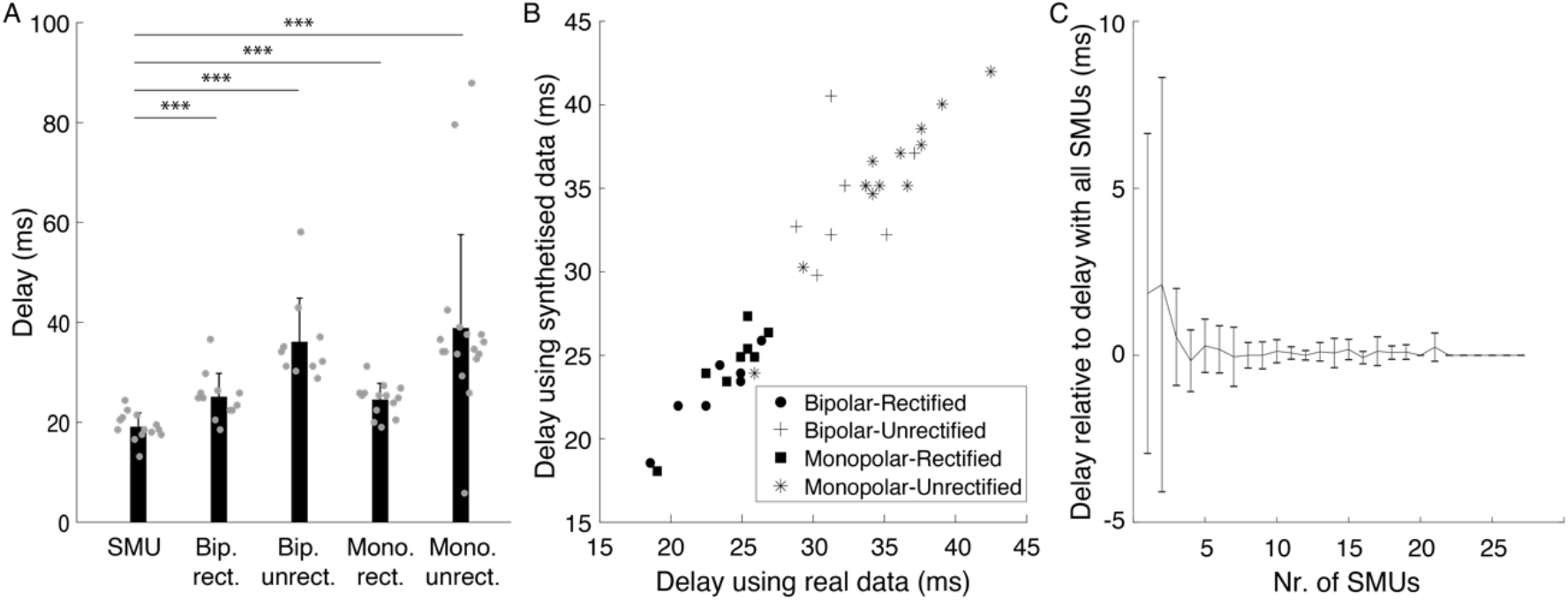
Comparison between EEG→SMU and EEG→EMG beta transmission delay estimates in Part 2. (A) Beta transmission delay estimates using SMUs activity and four different EMG montages: bipolar rectified (Bip. rect.); bipolar unrectified (Bip. unrect.); monopolar rectified (Mono. rect.); monopolar unrectified (Mono. unrect.). Grey circles represent individual data. *** p < 0.001. (B) Beta EEG→EMG delay estimates with real (X-axis) and synthetized EMG from the decomposed SMUs (Y-axis). (C) Group results of the EEG→SMU beta delay estimates obtained using increasing numbers of SMUs relative to estimates obtained considering all the decomposed SMUs in each subject.

To ensure that differences in the results with SMUs and EMG were solely due to the effect of the MUAPs, we synthetized EMG signals from the spiking SMU activity. Delay estimates with synthesized EMG were strongly correlated with delays obtained using real EMG data (R = 0.964; P < 0.01) (Fig. 4B). This high correlation indicates that differences observed between EEG→SMU and EEG→EMG delay estimates are mainly caused by the filtering effect of MUAPs. It also suggests that decomposed SMUs contain the vast majority of the relevant information in the EMG.

Finally, we also measured how EEG→SMU beta transmission delay estimates change with increasing samples of SMUs considered. This was done to assess if the activity extracted from the decomposed SMUs sufficiently describes the information of the entire pool of active motoneurons innervating the muscle. Results of this analysis show that stable EEG→SMUs delay estimates could be obtained in all subjects when activity of just four or more decomposed SMUs were combined (Fig. 4C). Therefore, the numbers of decomposed SMUs per subject obtained in this study should be sufficient to yield accurate delay estimates.

#### PART I-C. Integrated EEG gives better delay estimates than raw EEG

In the previous sections, EEG→SMU beta transmission delay estimates are obtained assuming that the descending beta activity does not undergo any phase delays between the point where EEG is recorded and the measurement point at the muscle. However, this is not correct, since pyramidal tract neuron spiking is related to the integral of nearby recordings of local field potential (Baker et al., 2003). This implies that EEG signals (recorded here with a polarity defined as negative upward) undergo a phase shift of π/2 in their integration by the pyramidal neurons. This phase shift must be taken into account as it affects beta transmission delay estimates obtained using the cumulant density function, which works in the time domain and is blind to phase delays.

Figure 5A illustrates the average cumulant between raw EEG and pooled SMU activity across subjects (grey trace). To correct for the phase shift between EEG and corticospinal neuron activity, we integrated the EEG by replacing each value with its cumulate sum up to that point; the cumulant determined with integrated EEG (∫ EEG dt) is illustrated with the black track in Fig. 5A. The delay estimated from the average cumulant peak shifted later, from 19.0 to 28.3 ms (dotted vertical lines).

**Figure 5.**
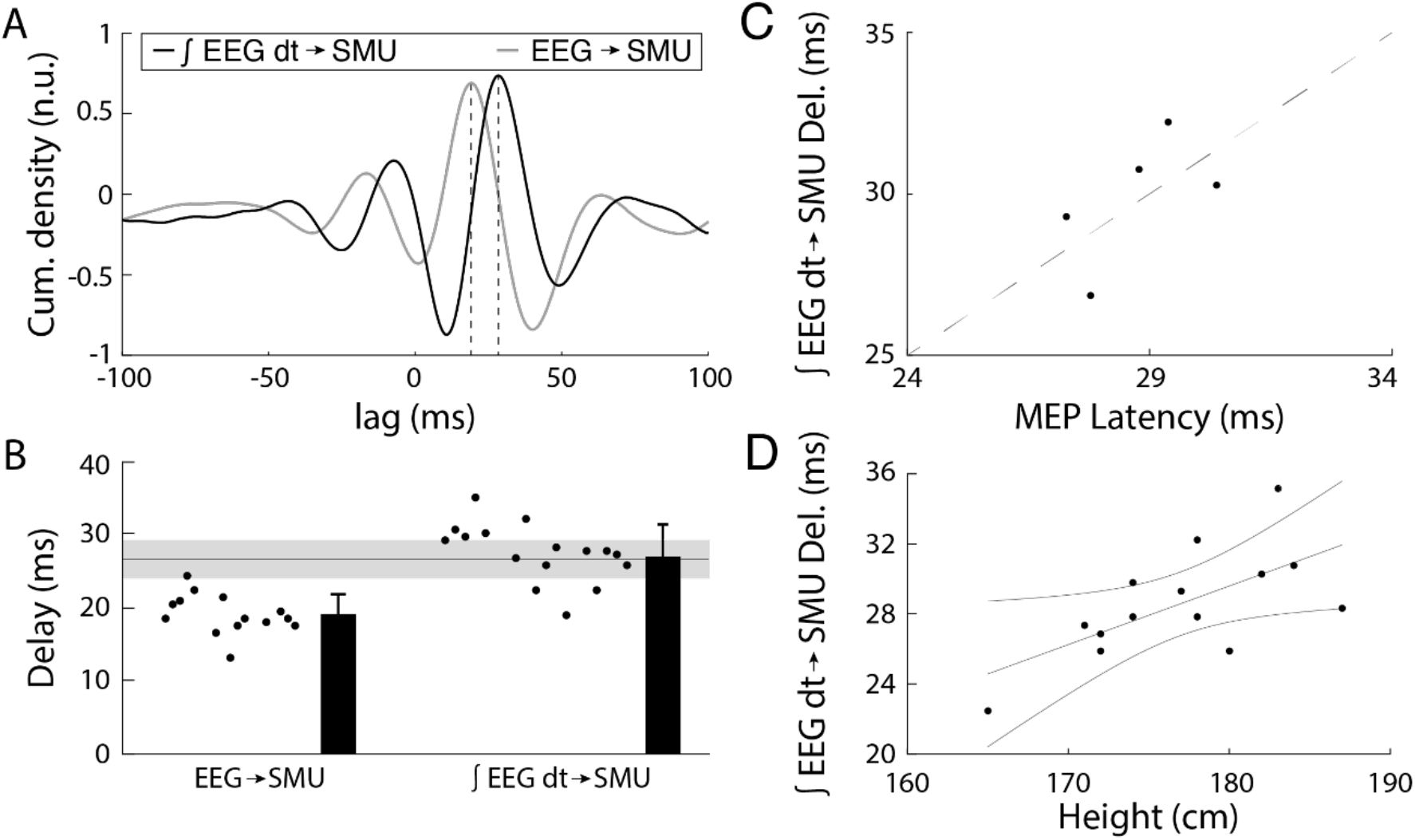
(A) Average cumulant densities obtained between the EEG and SMU activity (grey line) and between the integrated EEG (∫ EEG dt) and SMU activity (black line). (B) Beta transmission delay estimates when based on EEG without phase modifications (left) versus when considering that spike trains in pyramidal tract neurons relate to ∫ EEG dt (right). The horizontal solid line and grey area indicate the mean and SD of TA MEP latencies in previous studies using large sets (n = 89) of healthy individuals (Rossini et al., 1999). (C) ∫ EEG dt →SMU beta transmission delay estimates versus MEP latencies. A diagonal dashed line with a slope of 1 and intercept of 1 ms is included. (D) ∫ EEG dt→SMU beta transmission delay estimates versus subjects’ heights and linear fit with confident intervals at 95%.

Figure 5B presents individual and average results across subjects. By taking into account the phase shift caused by integrating EEG activity, the average cortical→SMU beta transmission delay became 28.6 ± 3.1 ms, which was 9.4 ± 1.0 ms longer than if the phase shift was not accounted for.

Compared to EEG→SMU delay estimates, Cortical→SMU delay estimates including the integration of EEG were much closer to previously reported measures of MEP latency for TA based on recordings from large samples of subjects (26.7 ± 2.6 ms; horizontal line and shading show mean ± SD in Fig. 5B) (Rossini et al., 1999). This is further confirmed when comparing estimated beta transmission delays with the MEP latencies measured in 5 subjects in our study (Fig. 5C): the difference between the two measures was 1.14 ± 1.60 ms. This suggests that beta is transmitted to the muscles through the fastest corticospinal axons. If Cortical→SMU beta coherence is determined by the most direct and fast descending projections (and not by varying groups of descending axons across individuals), assuming that conduction velocities of these fastest projections is comparable across subjects, then it is expected that beta transmission delays correlate with the travelled distances to the muscles (Jaiser et al., 2015). Fig. 5D shows the relation between Cortical→SMU beta transmission delay estimates and subjects’ heights. Delays correlated significantly with subjects’ heights (R = 0.651, P = 0.006).

### PART II – Corticomuscular transmission is determined by the fastest descending axons

Having removed methodological confounding factors from estimates of corticomuscular beta transmission delay, we found delays close to the fastest descending pathways as assessed by TMS. This is somewhat unexpected, given the large preponderance of slow fibers in the corticospinal tract. We explored how selective transmission by the fastest routes can be achieved in Experiment 3 by simulation. The distribution of axon diameters from a previous careful anatomical study in monkeys (Firmin et al., 2014) was modelled using a lognormal function, which provided a good fit (Fig. 6A, compare grey and black lines). The fitted lognormal function was then used to generate distributions of axon diameters, which were converted to conduction velocities assuming a Hursh factor of 6, and to transmission delays by assuming a given axon length. Figure 6B illustrates the distribution of transmission delays of the simulated axons for a distance of 800 mm, estimate to be close to the value for the human TA motoneuron pool (Jaiser et al., 2015). Using this distribution, we then simulated different scenarios in which beta activity was restricted to axons conducting faster than a certain threshold.

**Figure 6.**
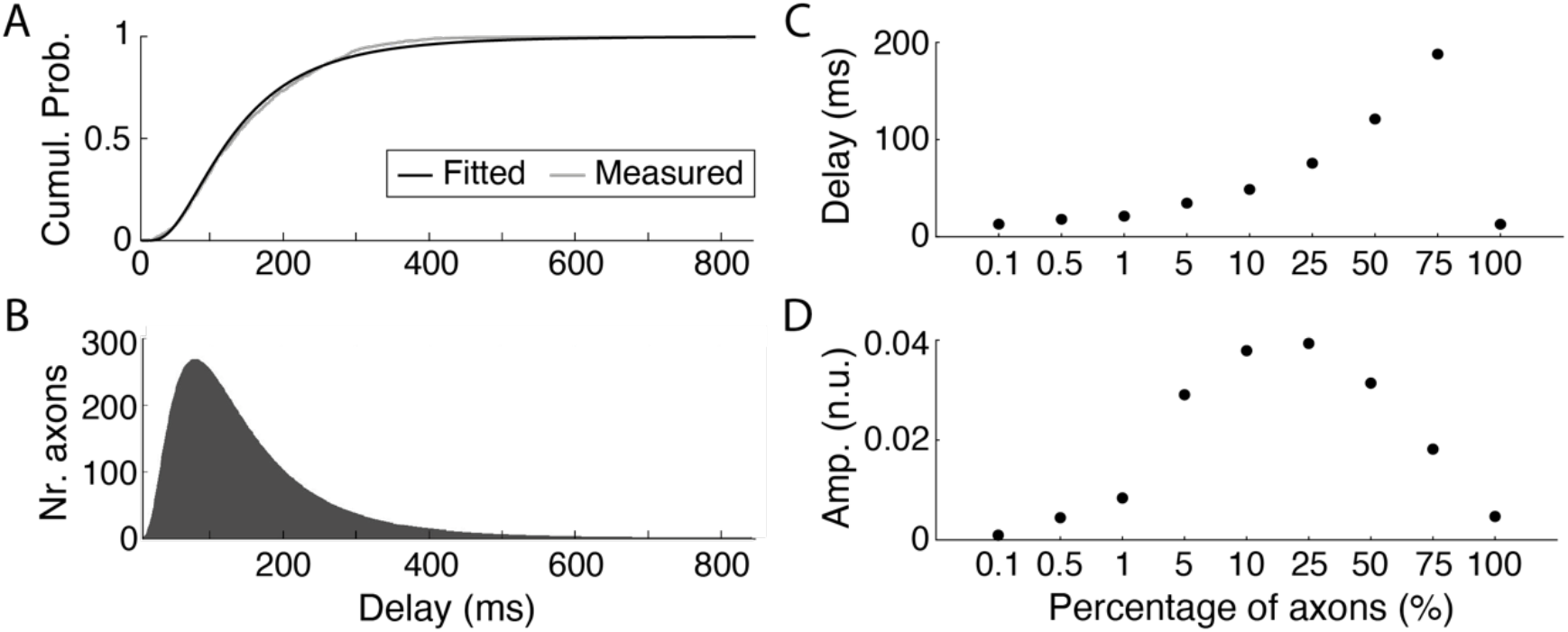
Results of the transmission and posterior summation of beta signals (band pass-filtered noise within 20-30 Hz) simulating the propagation of beta activity through the corticospinal tract to lower motoneurons. Simulations use a population of 10^6^ axons of 800 mm of length (similar to the estimated length of corticomotoneural cells to the TA) and axon diameters (and conduction velocities) based on previously published work (Firmin et al., 2014). (A) Cumulative probability function of delay estimates considering the measured axon diameters in Monkey CS28 in Firmin’s article (grey line) and the axon diameter distribution obtained by fitting a lognormal function to actual data (black line). (B) Histogram of the distribution of transmission delays used for the simulation (obtained using the fitted lognormal function shown on panel A). (C-D) Transmission delays and normalized rms amplitudes for different scenarios simulated in which different percentages of the fastest axons transmit the (cortical) beta signal. Amplitudes panel D result from summing the contributions of all axons transmitting beta and dividing by the total number of axons simulated (10^6^) and by the rms amplitude of the input signal.

Fig. 6C shows how the estimated transmission delay changed as the conduction velocity threshold altered. Unsurprisingly, when only the fastest 0.1% of the axons were allowed to transmit the beta oscillations, the measured conduction delay was short (13 ms), implying a conduction velocity of ~62 m/s. As more and more slow axons were included in the transmission of beta, the estimated delay rose. However, surprisingly when all axons were included, the delay dropped again to be the same as in the case where only the fastest 0.1% axons transmitted beta oscillations. This is because cancellation occurs over such a wide delay range: for a delayed version of the input oscillation, phase cancellation occurs due to the summation with other delayed versions of the same signal that arrive earlier and later to the end of the transmission channel. In this context, the only signals that are not cancelled are those transmitted at the fastest possible speeds. As a result, only the fastest axons (those not resulting in relevant phase shifts of the oscillations transmitted along the whole axon length) can transmit an uncancelled signal. Fig. 6D plots the corresponding normalized rms amplitude of the transmitted beta oscillation. Again, the amplitude rises with the inclusion of slower axons, but then falls as cancellation begins to influence the transmission. Note however that the amplitude when all axons contributed was 5×10^−3^ (normalized units), compared with 0.9 ×10^−3^ when only the fastest 0.1% of axons contributed.

Fig. 7 shows how transmission delays (Fig. 7A) and normalized rms amplitudes of the transmitted signals (Fig. 7B) change as a function of the length of the axons transporting beta. Results shown are for two extreme cases where either all axons or only the 0.1 % fastest axons transmit beta inputs. In the case where all axons transmit beta activity and contributions from slow axons are mutually cancelled, transmission delays remain relatively stable with small increases while amplitudes decay exponentially as axon lengths increase. This implies that longer transmission channels intrinsically lead to increasingly more selective transmissions involving only the fastest axons. By contrast, when only the 0.1 % fastest axons transmit beta activity, delays relative to the fastest axons increase linearly with the simulated axon lengths, which reflects a relatively stable conduction speed in this scenario. In this case, amplitudes remain relatively unchanged at very low levels since only a small number of axons contribute to beta transmission.

**Figure 7.**
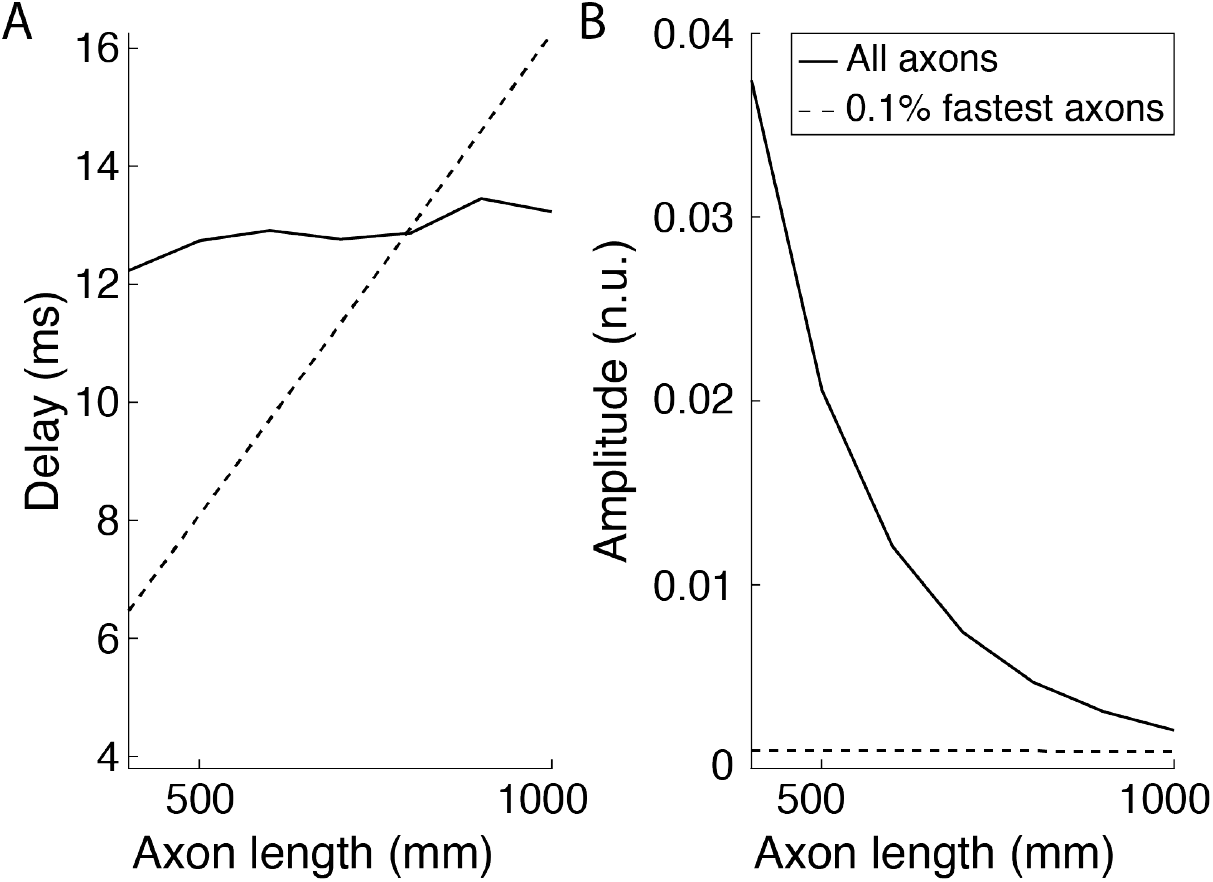
Changes as a function of the lengths of the axons of (A) beta transmission delays; (B) normalized rms amplitudes of the transmitted signals. These amplitudes resulted from summing the contributions of all axons transmitting beta and dividing by the total number of axons simulated (10^6^) and by the rms amplitude of the input signal. Results for two cases are plotted: all axons transmit beta activity (solid lines); only the 0.1 % fastest axons transmit beta activity (dashed lines).

## Discussion

Here we propose and validate methods to improve delay estimates in the corticomuscular transmission of beta rhythms. Applying these approaches to experimental data from human subjects revealed delays only a little greater than expected from the fastest corticospinal pathway. Most previous animal and human studies have worked exclusively on fast corticospinal transmission, due to unavoidable biases introduced by available recording and stimulation methods. Because much more is known about fast-conducting corticospinal components, it may be tempting to view our results as fitting comfortably with current knowledge. However, measuring propagation of endogenous beta oscillations is not inherently biased according to conduction velocity. Recent studies have emphasized that fast-conducting fibers make up only a tiny fraction of the corticospinal tract (Firmin et al., 2014; Kraskov et al., 2019, 2020). We might expect corticomuscular coherence delays to be much longer, perhaps close to the average conduction delay across all corticospinal fibers – and indeed long corticomuscular delays have been reported previously, but using approaches which we show here can yield overestimates (Witham et al., 2010, 2011). The finding that delays using improved methods actually match those from the fastest fibers is an unexpected result, which provides an important insight into motor control.

### Reliable estimates of cortical beta transmission delay to muscles

We found three factors which could lead to erroneous delay estimates in previous studies, and showed using simulated and experimental data how to compensate for them to produce reliable measures. The first is the nature of the analysis applied to extract delays in a bidirectional system, such as between the cortex and muscles (Riddle and Baker, 2005b; Baker, 2007). A common approach is to use directed coherence (Granger causality, (Granger, 1969)), and to measure the slope of the phase-frequency relationship (Witham et al., 2010, 2011). However, for bidirectional communication over a restricted bandwidth, delays are overestimated (Fig. 2). The peak time of the cumulant density yields unbiased delays, so long as the bandwidth is not very narrow (close to a pure sinusoid), a condition met by experimental data (Fig. 3).

When using directed coherence to estimate delays, constant phase shifts in one of the signals alter the offset of the regression line fitted to the phase-frequency relationship. This has no impact on delays, which are extracted from the regression slope. By contrast, a disadvantage of the cumulant density is that constant phase shifts change the peak time and hence the estimated delay. We know that corticospinal cells discharge with a ~90 degrees phase shift relative to local field potential oscillations (Baker et al., 2003); this phase shift can be corrected for by using the mathematical integral of the EEG (Baker et al., 2003).

A final relevant factor, often not considered previously, is the influence of the EMG. Surface EMG can be modelled using a cascade of multidimensional filters in time and space that replicate how the motor unit transforms motoneuron spiking to signals at the skin (Farina et al., 2002). The duration of the MUAP affects corticomuscular delay estimates (Williams and Baker, 2009). This can be avoided by decomposing EMG to SMUs, and instead using the pooled SMU spike train in analysis.

### Corticomuscular beta transmission is determined by the fastest corticospinal neurons

Our results indicate that corticomuscular beta transmission delays are 1-2 ms longer than the onset latencies of MEPs evoked by TMS. It is a familiar argument when studying stimulus-evoked electrophysiological responses that the earliest responses must reflect the action of the fastest conducting pathway, with the smallest number of interposed synapses (Witham et al., 2016). The close match of MEP and beta delays suggests that transmission is dominated by the fastest corticospinal neurons. The predominance of the fastest projections is unexpected, given that slowly-conducting neurons (<20 m/s) greatly outnumber fast ones (<3% of the population, (Firmin et al., 2014)). By simulating the summation of beta signals transmitted with transmission delays distributed according to the neuroanatomical data of Firmin et al., (2014), we conclude that there are two different scenarios which might explain the results.

In the first scenario, all corticospinal neurons transmit beta activity equally, but phase cancellation occurs between the contributions of slow corticospinal axons. This results from the characteristics of the delay distribution resulting from the axon diameter distribution (Fig. 6B). There is a sharp increase at short delays, and a long tail which declines gradually with long delays. Whilst consistent with the delay estimates, this scenario has associated implications. Phase cancellation will also occur for transmission of frequencies outside the beta band, but the extent depends on an interaction of the oscillation frequency, conduction velocity distribution and conduction distance. Supplementary Figs. 1–2 show comparable results to Figs. 6-7, but for a transmitted signal in the 8-12 Hz band. The effective transmission delays would be ~15 ms slower than where beta signals are transmitted. Even slower delays would occur for the frequencies below 5Hz which typically characterize voluntary movements (Churchland et al., 2012). It seems implausible that the motor system could operate effectively with such long and frequency-dependent delays, unless descending motor commands to muscles were modulated at higher frequencies (Watanabe and Kohn, 2015).

A second implication of this possibility is that the amplitude of beta activity reaching motoneurons should exponentially decay as the length of corticospinal axons increases (Fig. 7). This is at odds with empirical observations that corticomuscular coherence is larger in lower than upper limb muscles, although the central conduction distances are greater to lumbar than cervical spinal segments (Ushiyama et al., 2010).

An alternative scenario consistent with our results is that only a small proportion of corticospinal neurons with the fastest conduction velocity is effectively involved in transmitting beta activity. In this case, beta transmission delays would change proportionally to the axon lengths, and the beta amplitudes of motoneuron inputs would be determined by the number of axons involved, with little appreciable cancellation. This seems more plausible, and better fits experimental data. Such selective beta transmission could occur in two ways (which are not exclusive). Firstly, beta could only be carried down the corticospinal tract by fast axons. This would require that slow neurons are excluded from participating in the cortical circuitry which generates beta oscillations. It is known that fast corticospinal cells do not just read out the activity of a cortical oscillator (Baker et al., 2003), but are an integral part of the oscillatory generator (Jackson et al., 2002). It is possible that the intracortical connections of slowly-conducting corticospinal cells do not place them within this network. Even if connections are not so specifically organized, aspects of the single cell physiology might have a similar effect. Low neural firing rates reduce the efficiency with which beta oscillations are encoded in spike discharge (Baker et al., 2003; Lepage et al., 2011). Slowly conducting cells might have lower firing rates, although to date there is no data available to address this. In primate motor cortex, a wide range of pyramidal cells express the Kv3.1b channel (Soares et al., 2017). This could support more rapid repolarization after an action potential and hence higher firing rates, but does not seem to be restricted to only large cells which might contribute fast corticospinal axons.

Secondly, it is possible that, although slow corticospinal axons carry beta oscillations to the cord, they do not make connections capable of propagating this activity to motoneurons. Spike triggered averaging in awake monkey suggests that some corticospinal fibers with slower conduction velocities can make monosynaptic connections to motoneurons (down to an estimated 12 m/s, applying the conduction distances quoted in Firmin et al. (2014) and Kraskov et al. (2019) to antidromic latencies in (Lemon et al., 1986)), a conclusion which is supported by work in anesthetized animals (Witham et al., 2016). However, whether the very slowest fibers make such contacts, and whether these are as strong or as frequent as for the fast fibers, remains unknown. In the corpus callosum, larger diameter axons give off larger synaptic boutons (Innocenti and Caminiti, 2017), suggesting that they generate stronger inputs in their target neurons; it is not known whether this correlation also exists for cortico-motoneuronal connections. The situation regarding indirect, oligosynaptic influences on motoneurons, and the relative balance there between fast and slow corticospinal inputs, is similarly lacking in experimental data at present.

The observation that only the fastest corticospinal projections are responsible for corticomuscular beta coherence may allow a better understanding about how coherence is altered in certain conditions. For example, in humans with a stroke or with motor neuron disease, beta corticomuscular coherence is reduced (Fisher et al., 2012; von Carlowitz-Ghori et al., 2014; Proudfoot et al., 2018). This may be because large corticospinal neurons are most vulnerable in these pathologies, and their loss leads to less reliable cortical transmission to muscles (Brownell et al., 1970; Kraskov et al., 2019).

### Limitations

One limitation of the present study is that our experimental validation used the tibialis anterior muscle. This is well suited to EMG decomposition (Del Vecchio et al., 2020), and is characterized by receiving a large amount of direct corticospinal contributions (Ushiyama et al., 2010). The generalization of our observations to other muscles with different proportions of efferent and afferent contributions is unproven here and should be tested in future works.

## ABBREVIATIONS

SMU: single motor unit
HD-EMG: high-density electromyography
EEG: electroencephalography
TA: tibialis anterior muscle
MUAP: motor unit action potential
MEP: motor evoked potential
TMS: transcranial magnetic stimulation

## Acknowledgments

We thank Roger Lemon for providing the raw data on axon diameters presented in Firmin et al (2014) and used here to model the distribution of conduction delays in the descending cortico-motoneural cells. We thank Paul Hammond for building the experimental platform used to record forces generated by the tibialis anterior muscle. We thank David Halliday for his comments and feedback on the use of Neurospec.

## Supplementary material

**Supp. Figure 1.**
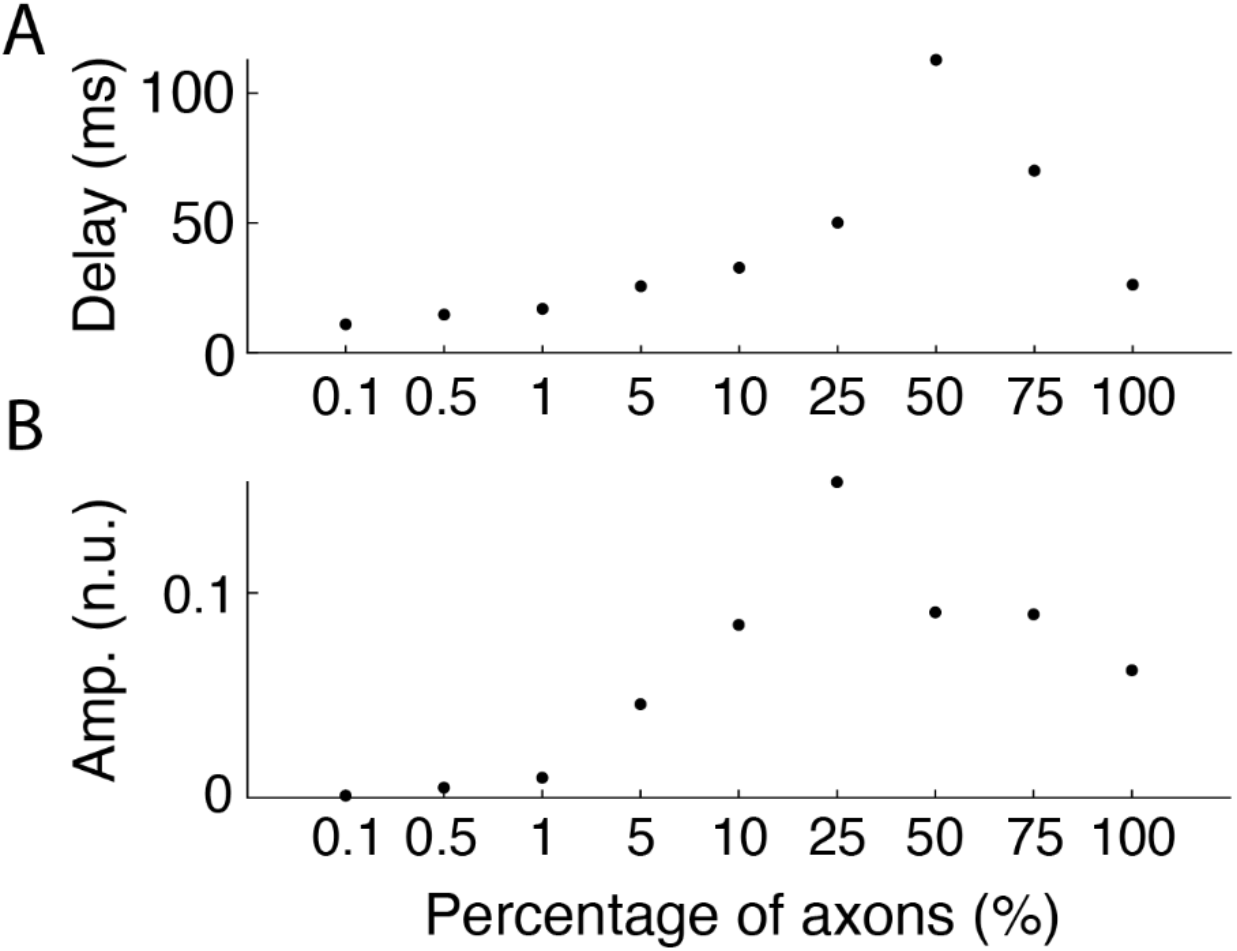
Results of the transmission and posterior summation of band pass-filtered noise within 8-12 Hz, simulating the propagation of alpha activity through the corticospinal tract to lower motoneurons. Simulations use a population of 10^6^ axons of 800 mm of length (similar to the estimated length of corticomotoneural cells to the TA) and axon diameters (and conduction velocities) based on previously published work Firmin 2014 (A-B) Transmission delays and amplitudes (in normalized units) of the transmitted signals for different scenarios simulated in which different percentages of the fastest axons transmit the (cortical) alpha signal.

**Supp. Figure 2.**
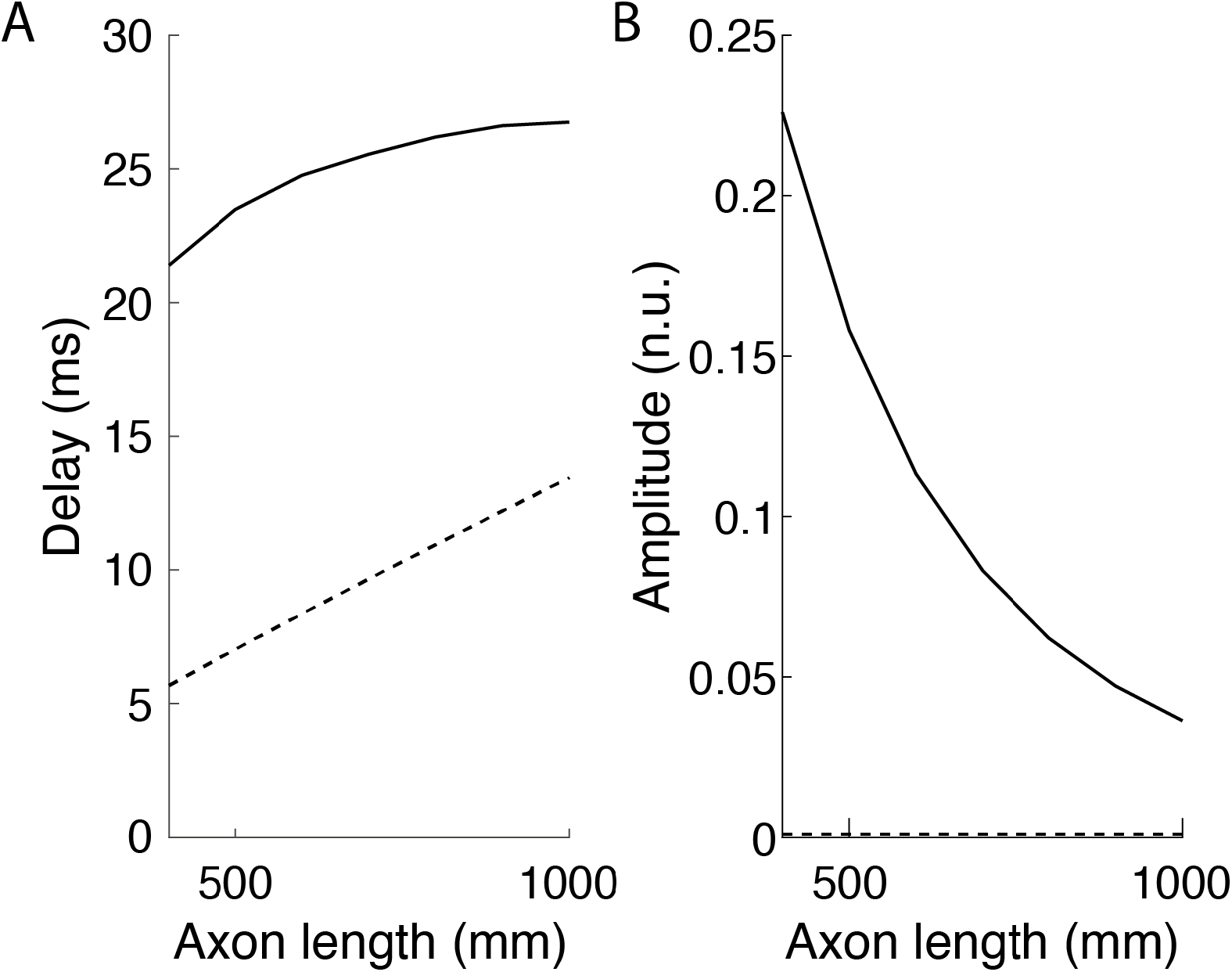
Changes with axon lengths of (A) alpha (white gaussian noise filtered between 8-12 Hz) transmission delays; (B) amplitudes (in normalized units) of the transmitted signals resulting from summing all axons transmitting alpha activity. Results for two cases are plotted: all axons transmit alpha activity (solid lines); only the 0.1 % fastest axons transmit alpha activity (dashed lines).

## Notes

*CONFLICT OF INTEREST:* The authors declare no competing financial interests.

*FUNDING:* This study was funded by the European Research Council (Synergy Grant Natural BionicS, contract #810346).

### Competing Interest Statement

The authors have declared no competing interest.

